# Pre-Sequencing Assessment of RNA-Seq Library Quality Using Real-Time qPCR

**DOI:** 10.1101/2025.07.08.663710

**Authors:** Kavya Kottapalli, Hsu Chao, Qi Jiang, Jimmonique Donelson, Nathanael Emerick, Sravya V Bhamidipati, Zeineen Momin, Mathew Tittu, Ziad M. Khan, Korchina Viktoriya, Marie-Claude Gingras, Donna Muzny, Richard Gibbs, Harsha Doddapaneni

**Affiliations:** Human Genome Sequencing Center, Baylor College of Medicine, Houston, TX, 77030, USA

**Keywords:** RNA-Seq, 18S qPCR, rRNA removal, depletion efficiency, oligo dT beads, Illumina, Poly A+, Total RNA

## Abstract

RNA sequencing (RNA-Seq) is an essential sequencing assay for studying transcriptome profiling. Ribosomal RNA (rRNA) comprises more than 80 - 90% of total cellular RNA, efficient rRNA removal is essential for accurately capturing the transcriptome, particularly to sequence low-abundance mRNAs. Inefficient rRNA removal during library preparation can result from variations in sample quality, preparation methods and handling. Estimating rRNA content in RNA-Seq libraries pre-sequencing is therefore challenging due to the absence of a reliable and cost-effective assessment method. This study addresses the issue by introducing a scalable qPCR-based assay targeting 18S rRNA to evaluate rRNA depletion efficiency pre-sequencing. qPCR efficiency was optimized using serial dilutions of Universal Human Reference (UHR) control and Ct thresholds were established using pilot data from 644 libraries. Following this optimization, analysis of 1,748 Total RNA-Seq libraries and 445 Poly A+ two widely used RNA-Seq library methods, demonstrated a strong correlation between 18S rRNA qPCR results and post-sequencing rRNA rates. This assay was also used to evaluate the performance of Oligo dT beads from four different vendors to enrich mRNA. This 18S rRNA qPCR assay is a cost-effective, scalable approach for reliably predicting rRNA read percentage in RNA-Seq libraries pre-sequencing.

**Method Summary:** The 18S rRNA-specific qPCR assay is a cost-effective method for evaluating rRNA depletion efficiency in RNA-Seq libraries prior to sequencing by targeting the 18S ribosomal RNA. qPCR efficiency was optimized, and Ct thresholds were set from pilot data and validated across 1,748 Total RNA and 445 Poly A+ RNA-Seq libraries. A Ct threshold of ≥16 for Total RNA and Ct 13-14 or greater for Poly A+ libraries demonstrates a strong correlation between qPCR results and post-sequencing rRNA read percentages. The assay enables early identification of poorly depleted samples, reducing unnecessary sequencing costs. Additionally, the method was employed to benchmark oligo-dT beads from four vendors, showing its utility in evaluating RNA-seq library preparation kits. This scalable approach supports quality control in both research and high-throughput sequencing environments.

## 1. Introduction

RNA sequencing (RNA-Seq) has revolutionized transcriptome studies, enabling detailed analysis of gene expression, alternative splicing, and non-coding RNA [1–4]. High-throughput RNA-Seq, powered by massive parallel sequencing, has advanced molecular biology by uncovering genomic functions and identifying expressed genetic variants, including mutations and germline variations [5]. In cancer research, RNA-Seq plays a pivotal role for biomarker discovery, understanding drug resistance, cancer heterogeneity, evolution, and advancing immunotherapy [6].

RNA-Seq is resource-intensive, making it important to reduce reads from abundant transcripts like rRNA. A key challenge is the presence of ribosomal RNA (rRNA), which comprises more than 80-90% of Total RNA [7, 8], and if not efficiently removed, will pose a significant challenge in RNAseq experiments [2] Several strategies are employed for the removal of rRNA, including enzymatic or probe-based rRNA depletion, target enrichment or hybridization techniques, and the Poly A+ selection using oligonucleotide beads. These methods can significantly reduce rRNA content [7, 9–13]. Among these, Total RNA-Seq and Poly A+ RNA-Seq are the two most used methods for RNA library preparation. While effective under optimal conditions, rRNA removal methods vary with handling, sample quality, and instrumentation [7, 11, 14]. Without a thorough assessment, library preparation may yield high rRNA rates, increasing sequencing costs [14].

Therefore, there is a need for a methodology to assess rRNA removal efficiency pre-sequencing. Here, we present a cost-effective methodology of real-time quantitative polymerase chain reaction (qPCR) that can be used prior to sequencing to assess the efficiency of rRNA depletion in most used RNA-seq library preparation methods.

## 2. Materials and Methods

### 2.1. Sample QC

For this study, the Total RNA was analyzed using RNA 6000 Pico ( Agilent, Cat# 5067-1513) or Nano kits ( Agilent, Cat# 5067-1511) on an Agilent Bioanalyzer (2100) to determine the sample quality using RNA integrity number (RIN) and DV200% metrics. RNA samples used in this study were extracted from blood, tumor organoids and cell lines from various research studies processed in our laboratory. A total of 1,748 RNA samples were used to prepare Total RNA libraries, and 445 RNA samples were used to prepare Poly A+ libraries.

### 2.2. cDNA synthesis and Library Preparation

#### 2.2.1. Total RNA-Seq libraries

Whole transcriptome sequencing (Total RNA-Seq) data was generated using the Illumina TruSeq Stranded Total RNA LP Globin kit (Illumina, Cat # 20020613). To monitor sample and process consistency, 1 µl of the 1:50 diluted synthetic RNA designed by the External RNA Controls Consortium (ERCC) (4456740, Thermo Fisher) was added to 1 µg of Total RNA. Additionally, the Universal Human Reference RNA (UHR) (Cat# 750500, Agilent) was processed in parallel with the RNA samples as a process control.

Library preparation followed the Illumina TruSeq Stranded Total RNA Sample Preparation Guide; (RS-122-9007 DOC, Part # 15031048 Rev. E, October 2013) with a few modifications noted below. The rRNA and Globin depletion were performed simultaneously by mixing the RNA sample, binding buffer, and rRNA and Globin removal mix, followed by incubation at 68°C for 10 min and then at room temperature for 5 min. For purification, the sample was transferred to a tube containing rRNA/Globin Removal Beads. The sample and beads were thoroughly mixed by repeated pipetting and incubated at 50°C for 5 min followed by magnetic separation of the beads and collection of supernatant. This process was repeated 3 times to completely remove the rRNA and Globin Removal Beads. The rRNA and globin-depleted RNA sample was purified with Agencourt RNAClean XP beads (Beckman Coulter, Cat #. A63987) and eluted in 9 µl Elution buffer. To fragment the RNA, the eluate was combined with Fragmentation High mix and was incubated at 94°C for 3 min on a thermocycler (PTC-200, MJ Research, Inc.). The fragmented RNA was rapidly cooled on ice and used for first and second-strand cDNA synthesis as described in the preparation guide. The synthesized double stranded cDNA was eluted into 16.5 µl of the resuspension buffer.

Paired-end libraries were prepared on Beckman BioMek FXp liquid handlers using TruSeq kit reagents. The cDNA was A-tailed, followed by ligation of the TruSeq UD Indexes (Cat # 20022370), and amplified for 15 PCR cycles according to the manufacturer’s recommendation. AMPure XP beads (A63882, Beckman Coulter) were used for library purification. Libraries were quantified using a Fragment Analyzer (Agilent Technologies, Inc) electrophoresis system (Figure 1).

**Figure 1:**
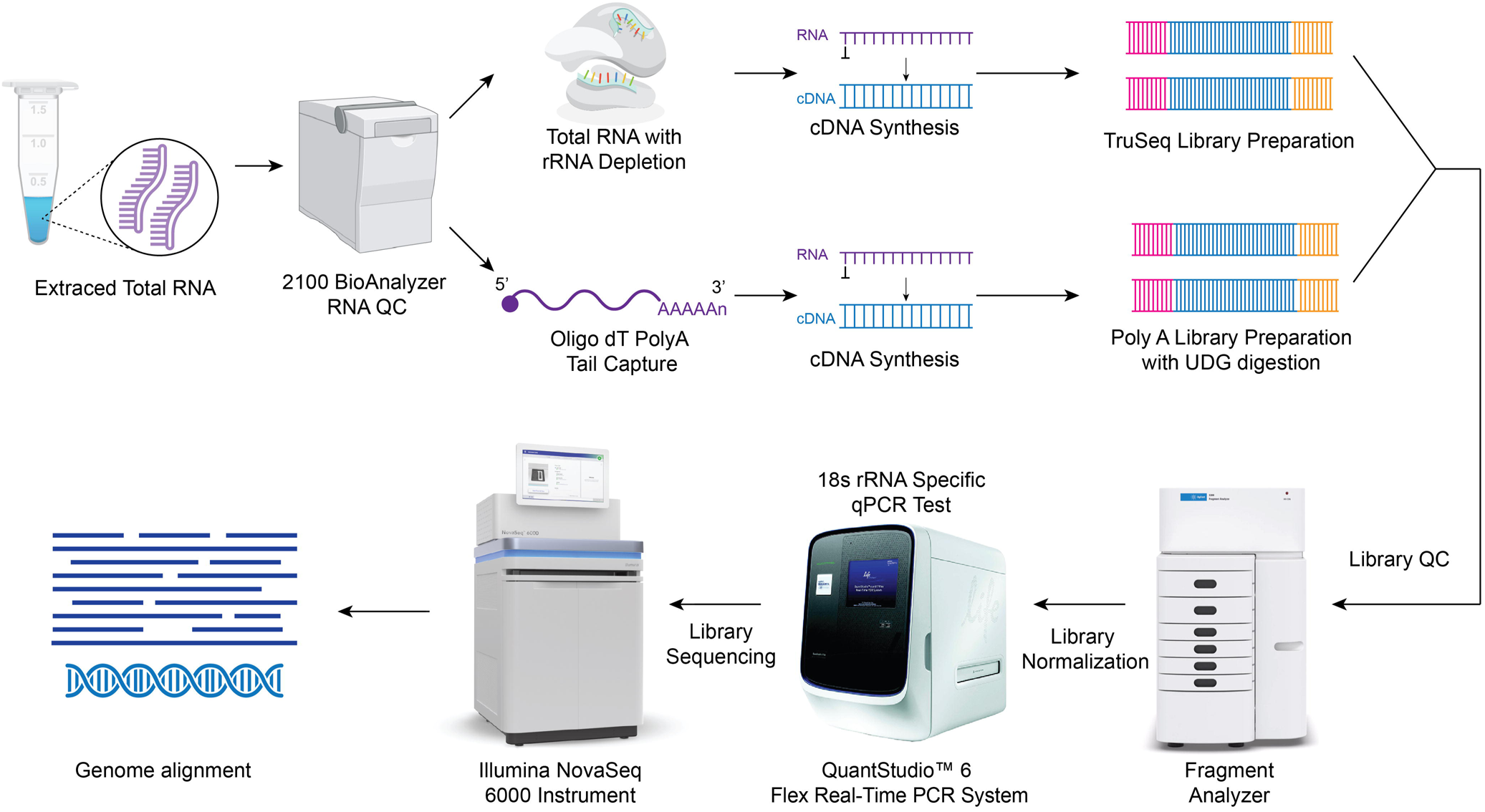
Schematic workflow: The workflow illustrates the various steps involved in library preparation and sequencing methodology. First row- RNA was received and quantified using Bioanalyzer and RNA was converted to cDNA. Second row– cDNA was used to generate Illumina libraries with molecular barcodes and these libraries were normalized to equimolar concentration and 18S rRNA qPCR was performed to obtain Ct values. Libraries above the established Ct thresholds were then sequenced on the Illumina NovaSeq 6000 instrument to generate 2x150 bp length reads.

#### 2.2.2. Poly A+ libraries

RNA sequencing (RNA-Seq) was performed using Total RNA extracted from fresh-frozen tissue to prepare strand-specific, Poly A+ RNA-Seq libraries for sequencing on the Illumina platform. On each plate, along with the samples, an aliquot of Universal Human Reference (UHR) RNA sample (Cat# 750500, Agilent) was processed, serving as a positive control for cDNA synthesis and library preparation. Briefly, Poly A+ mRNA was extracted from 1 μg Total RNA in 50ul using Oligo dT 25 Dynabeads (Life Technologies, Cat #. 61002) followed by fragmentation of the mRNA by heat at 94°C for 5 minutes (for the samples with RIN above 6) or less accordingly (for samples with RIN lower than 6). First-strand cDNA was generated utilizing NEB Next RNA First Strand Synthesis Module (E7525L; New England Biolabs Inc.) and NEB Next Ultra II Directional RNA Second Strand Synthesis Module (E7550L; New England Biolabs Inc.). The double strand (ds) cDNA was purified using 1.8X volume of AMPure XP beads (A63882, Beckman) and eluted into 23 μl 10 mM Tris buffer (Cat#A33566, Thermo Fisher Scientific). Oligo dT beads from four vendors (Vazyme, Qiagen, NEB, Thermo Fisher) were assessed according to the manufacturer’s protocols, and cDNA conversion was performed as mentioned above.

Libraries were prepared using Beckman BioMek FXp robots. Automated processes were employed for cDNA end repair, 3’-adenylation, ligation to Illumina multiplexing PE adaptors, UDG digestion and ligation-mediated PCR (LM-PCR). Libraries were amplified for 13 PCR cycles in 50 μl reactions containing 150 pmol of P1.1(5’-AAT GAT ACG GCG ACC ACC GAGA) and P3 (5’-CAA GCA GAA GAC GGC ATA CGA GA) primers and Kapa HiFi Hot Start Library Amplification kit (Cat# KK2612, Roche Sequencing and Life Science). A set of 96 unique dual-index barcodes (Illumina TruSeq UD Indexes, # 20022370) were used for sample barcoding. Libraries were purified with Agencourt AMPure XP beads (Beckman Coulter, Cat # A63882) after each enzymatic reaction and PCR amplification, and were quantified using the Fragment Analyzer (Agilent Technologies, Inc.) electrophoresis system (Figure 1).

#### 2.2.3. Sequencing on Illumina Novaseq6000

Sequencing was performed on the NovaSeq 6000 instrument using the S4 reagent kit (300 cycles) to generate 2x150bp paired-end reads. To achieve a sequencing depth of 50 to 100 million read-pairs per sample, between 35 and 70 RNA-Seq libraries were pooled per lane and loaded at 280 pM onto an S4 flow cell. Each lane included a 1% PhiX spike-in as an internal control to assess run quality.

### 2.3. Ribosomal RNA(18S) qPCR Assay

For rRNA qPCR, the 18S rRNA specific forward(F) primer F1565 (5’-CAGCCACCCGAGATTGAGCA-3’) and reverse(R) primer R1816 (5’-TAGTAGCGACGGGCGGTGTG-3’) [15] were custom synthesized by Integrated DNA Technologies (IDT) at a concentration of 100 µM. KAPA SYBR FAST qPCR Master Mix (2X) Universal (KK4601, 07959389001) was used and 5 µl of 100µM forward and reverse primers were added to 5 mL of qPCR master mix. Libraries were diluted to 10nM concentration using an Elution buffer based on the Fragment Analyzer (Agilent Technologies, Inc) concentrations. Libraries were further diluted 100-fold using water. The qPCR reaction was set up by adding 4 ul of 0.1 nM diluted library and 6 ul of 18S rRNA qPCR master mix with a final volume of 10µl. Amplification was performed on the Quant studio 6 Flex Real-Time PCR System (Thermo Fisher, USA). Using the following program (3 min at 95°C), followed by a PCR stage of 40 cycles (5s at 95°C, 30s at 60°C), melt curve stage (5s at 95°C, 30s at 60°C and 35s at 95°C). Serial dilutions of Total and Poly A+ mRNA UHR control libraries, previously sequenced with a known rRNA rate were used to optimize the library input amount for this qPCR reaction. The slope of the standard curve was used to determine the efficiency and exponential amplification of the qPCR reaction.

### 2.4 Data Analysis

The RNA-Seq analysis pipeline processes raw RNA sequencing data (FASTQs), performing thorough quality control and allowing alignment to either the GRCh37 or GRCh38 reference genomes (excluding alternate contigs). It incorporates software tools including, STAR v2.7.3a [16] for sequence alignment, Picard v2.22.5 for marking duplicate reads, and Sam tools v1.9 for BAM to FASTQ conversion. Additionally, RSEM v1.3.3 [17] is utilized for gene expression quantification, while RNA-SeqC v1.1.9 [18], Qualimap2 v2.2.1 [19], and ERCCQC v1.0 generate quality metrics. The pipeline employs FeatureCounts v2.0.1 to derive raw gene feature counts, facilitating subsequent analysis for differential gene and isoform expression using DESeq2 or edgeR packages.

## 3. Results

### 3.1 Human 18S rRNA qPCR optimization

In this study, a real-time qPCR assay was optimized as a cost-effective and high-throughput method for evaluating rRNA abundance, ensuring accurate and reproducible results. To test the efficiency of the qPCR reaction, serial dilutions (1:10) of Total RNA and Poly A+ UHR control libraries previously sequenced with known rRNA rates of 13% and 15% respectively, were used. Reactions were set up along with No template Control (NTC). Figure 2 below shows the PCR efficiency of the Total RNA and mRNA libraries. There is a two Ct Value difference between the two library types, which exceeds the minor difference in the rRNA reads percentages (Figures 2a and 2d).

**Figure 2:**
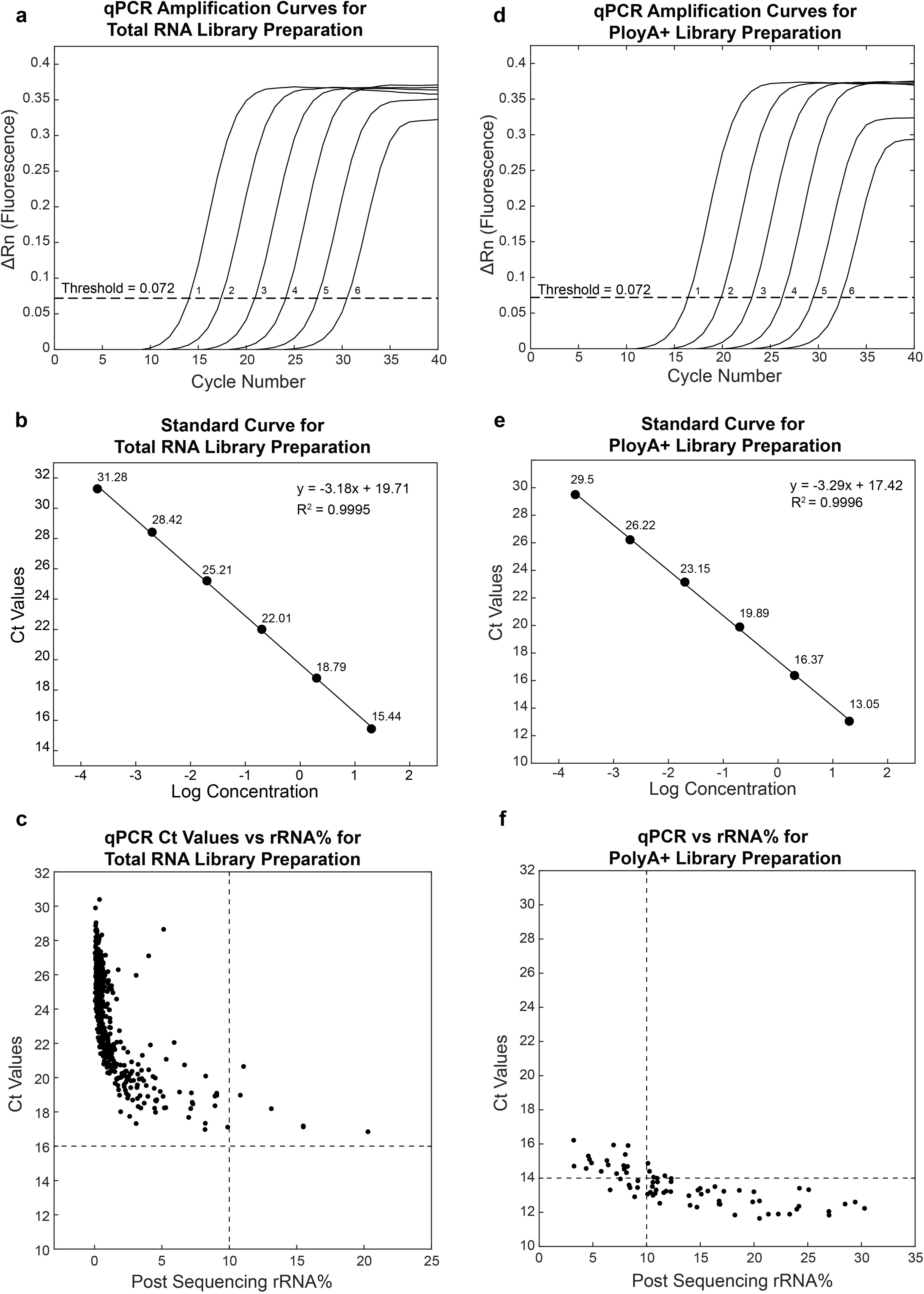
Total RNA library panels (a, b, and c) and Poly A+ library panels (d, e, and f) ; (a) Assessment of PCR efficiency using serial dilutions of the Total RNA UHR library control. (b) showing slope of -3.18, intercept of 19.7, and R-square value of 0.99 and Ct difference for each dilution of Total RNA libraries.(c) Pilot set of 570 Total RNA libraries. (d) Assessment of PCR efficiency using serial dilutions of the Poly A+ RNA UHR control and (e) showing slope of -3.29, intercept of 17.4 and R-square value of 0.99 and Ct difference for each dilution of Poly A+ libraries. (f) Pilot set of 74 Poly A+ RNA libraries.

A pilot set of 570 total RNA libraries was initially evaluated to determine the correlation between Ct values and rRNA read percentages after sequencing (Figure 2C). Based on these findings, a Ct threshold of 16 was established as optimal for screening total RNA samples. A similar approach was taken to correlate Ct value with post sequencing rRNA read percentages of 74 Poly A+ libraries and the data there suggested that a Ct threshold of 16 was stringent and would exclude 73/74 of libraries (Figure 2F). Therefore, a lower threshold of Ct 14 which is the same 2 Ct cycle offset as observed in Figures 2a and 2d between the two library types, was used to screen Poly A+ libraries.

### 3.2 Use in High-throughput production

#### 3.2.1 Total RNAseq

Following validation of the 18S qPCR assay, the method was implemented in production to screen 1,748 samples extracted from whole blood and flash-frozen tissues. Using a Ct value threshold of 16, 64 Libraries (3.5%) were excluded, and the remaining libraries were sequenced. Of the 1,684 libraries with Ct value above 16, 1,511 libraries were found to contain less than 1% rRNA read percentages (Figure 3) . The Ct value of these libraries varied from 20-30. For the remaining 172 libraries, which had rRNA read percentages between 1–10%, the corresponding Ct values ranged from 16 to 27. Overall, 99.6% of the libraries with Ct value above 16 have rRNA read percentages of less than 10% and the remaining 0.4% libraries were outliers.

**Figure 3:**
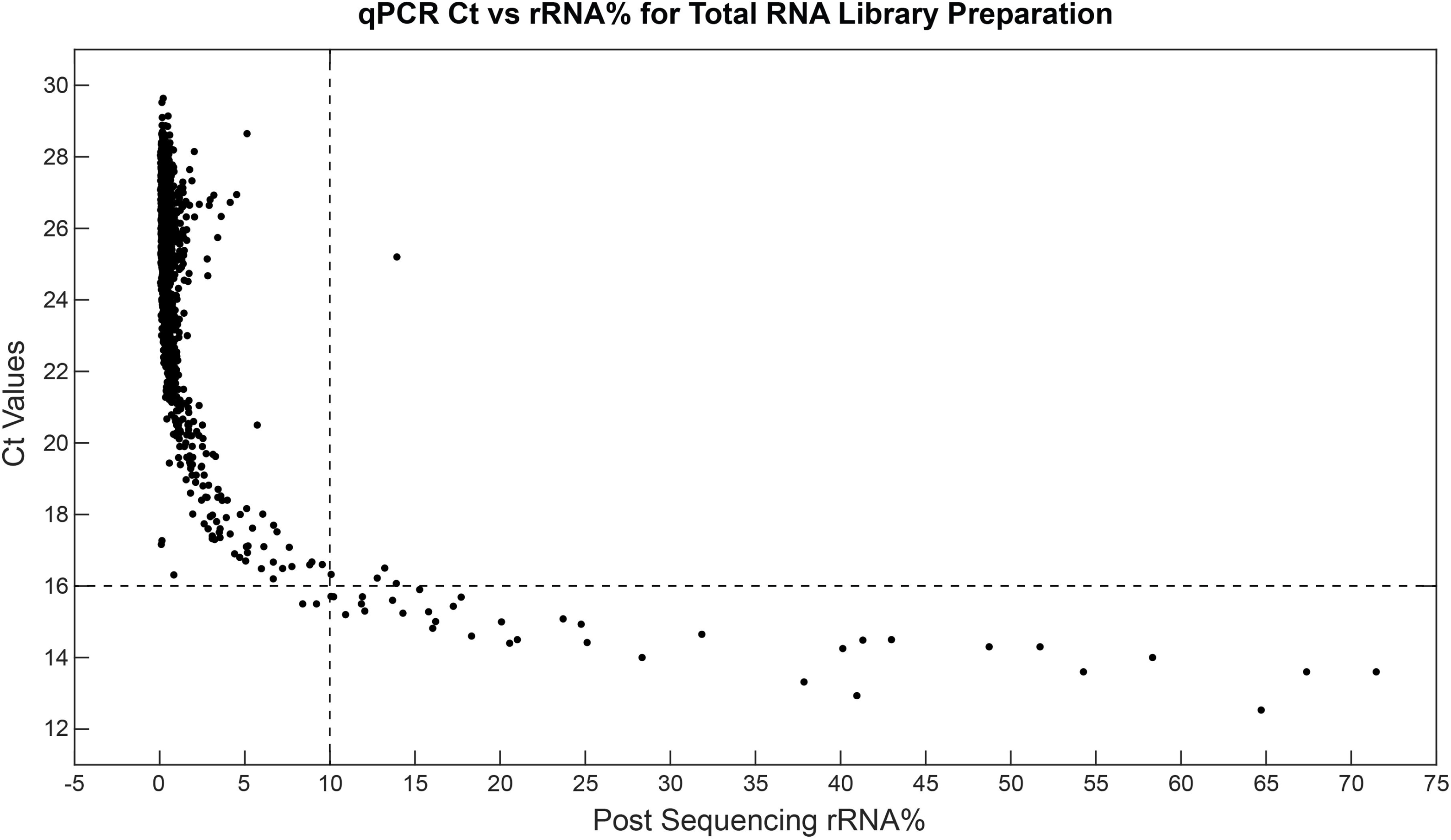
qPCR Ct values versus post sequencing rRNA read percentages for 1748 Total RNAseq Libraries. Ct value of 16 and post sequencing rRNA read percentage of 10 are shown in dotted lines.

Due to the lack of additional starting RNA samples, 36 out of 64 of the libraries that were initially rejected were also sequenced. All these libraries exhibited elevated rRNA read percentages, ranging from 10.9% to 76%. (Figure 3). To compensate for the rRNA reads in them, additional reads were generated.

#### 3.2.2 Poly A+ RNAseq

This method was implemented in production to evaluate 445 Poly(A)+ RNA-seq libraries generated from flash-frozen tumor tissues and cell lines (Figure 4). Using the Ct value threshold of 14, 292 libraries were passed for sequencing. Among these, 178 Libraries with Ct values between 16 and 29 showed low rRNA content (0–8%). There were 53 libraries with Ct values between 15 and 16, with rRNA read percentage ranging from 1% –8%. Additionally, 36 Libraries with Ct values between 14 and 15 had an rRNA read percentage ranging from 1% – 9%. Among the 292 libraries with Ct values ≥14, 25 showed post-sequencing rRNA content between 10% – 18% and were considered outliers.

**Figure 4:**
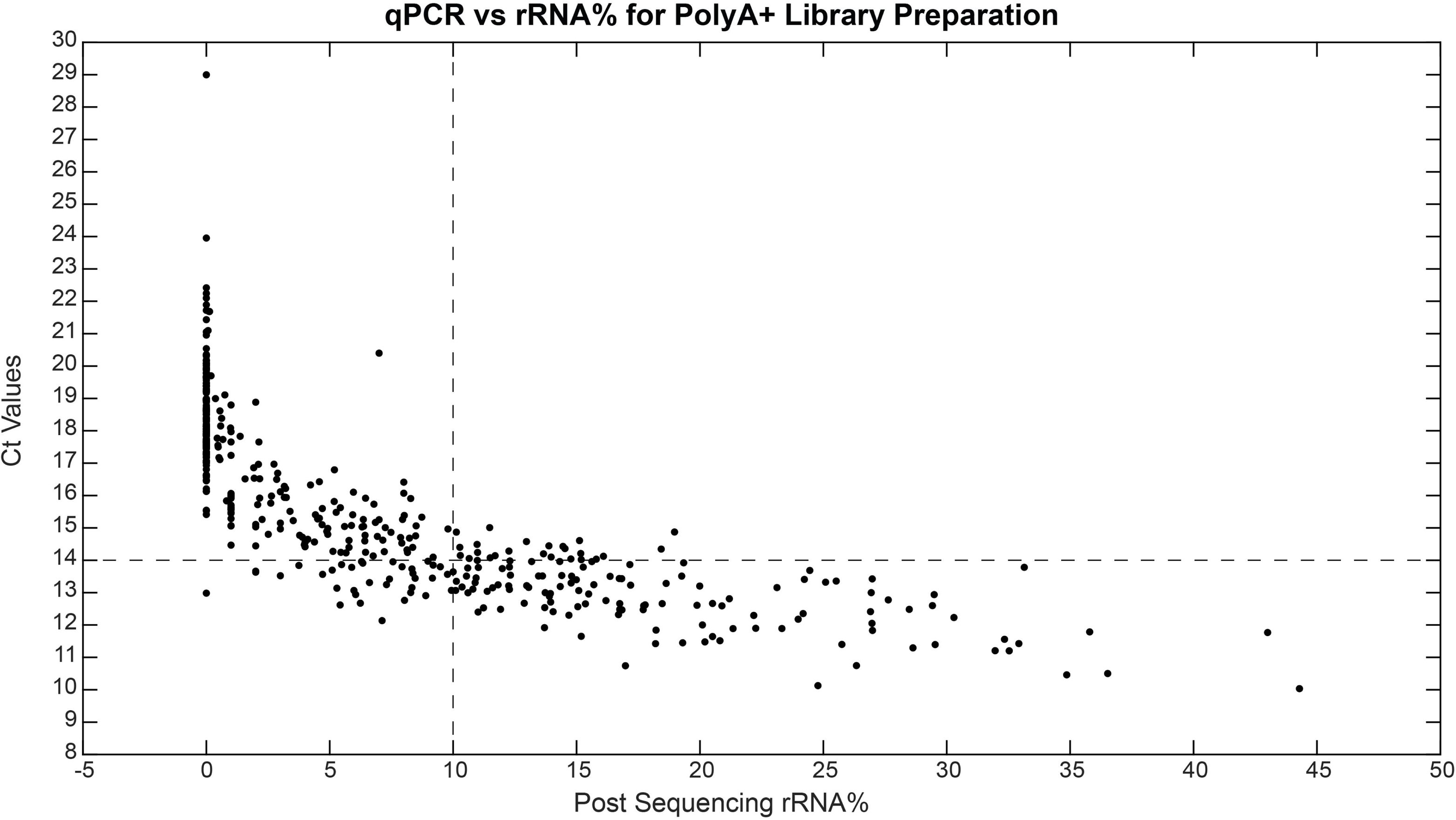
qPCR Ct values versus post sequencing rRNA read percentages for 445 Poly A+ RNAseq Libraries. Ct value of 14 and post sequencing rRNA read percentage of 10 are shown in dotted lines.

The remaining 153 libraries, which had Ct values below 14, were initially rejected due to expected high rRNA levels. However, due to limited sample availability, all these were sequenced, of which 118 libraries showed rRNA read percentage as expected, an elevated percentage ranging from 10% – 44%. The remaining 35 libraries showed an average of 6.9% rRNA reads. Interestingly, 29/35 libraries have a Ct value between 13 and 14 suggesting that libraries in this Ct range may still yield acceptable rRNA levels.

### 3.3 rRNA qPCR utilization in new RNAseq kit testing

This method was employed to assess mRNA enrichment using Oligo dT beads from four different vendors: New England Biolabs (NEB), Vazyme Biotech, Thermo Fisher Scientific, and Qiagen (Table 1).

**Table 1:** qPCR Ct values and post sequencing results for Oligo dT bead testing from four vendors using UHR control samples.

| Vendor | rRNA qPCR Results (Ct) | Reads (M) / Sample | Read Mapping Rate (%) | Unique Rate of Mapped (%) | Duplication Rate (%) | Intergenic Rate (%) | Intragenic Rate (%) | Exonic Rate (%) | Intronic Rate (%) | Genes Detected | Transcripts Detected | rRNA Rate(%) | Strand Specificity (%) |
| --- | --- | --- | --- | --- | --- | --- | --- | --- | --- | --- | --- | --- | --- |
| NEB | 16.57 | 191.31 | 96.97 | 66.01 | 33.99 | 2.42 | 97.51 | 85.43 | 12.08 | 31502 | 175822 | 1.45 | 97.98 |
|  | 16.93 | 146.63 | 96.87 | 69.44 | 30.56 | 2.42 | 97.52 | 85.53 | 11.99 | 30384 | 173117 | 1.41 | 97.95 |
| Vazyme | 20.70 | 91.67 | 97.64 | 71.67 | 28.33 | 1.35 | 98.58 | 90.79 | 7.79 | 27711 | 166490 | 0.15 | 98.37 |
|  | 21.21 | 149.73 | 97.60 | 66.37 | 33.63 | 1.36 | 98.57 | 90.77 | 7.80 | 29496 | 171775 | 0.15 | 98.50 |
| Thermo | 12.47 | 187.85 | 96.81 | 61.91 | 38.09 | 7.68 | 92.26 | 75.78 | 16.48 | 31262 | 174981 | 12.29 | 96.16 |
|  | 13.19 | 126.82 | 96.49 | 65.41 | 34.59 | 7.07 | 92.87 | 76.89 | 15.98 | 29490 | 170382 | 11.13 | 96.62 |
| Qiagen | 19.20 | 193.25 | 97.49 | 67.59 | 32.41 | 1.67 | 98.25 | 88.19 | 10.06 | 30904 | 175775 | 0.61 | 98.18 |
|  | 19.08 | 170.19 | 97.46 | 70.61 | 29.39 | 1.66 | 98.26 | 88.37 | 9.89 | 30366 | 174497 | 0.63 | 98.10 |

Libraries were prepared in duplicates for each bead type, followed by rRNA qPCR analysis. The Ct values in the libraries ranged from 12 to 21, indicating varying levels of rRNA percentages (Table 1). Vazyme’ s oligo dT beads showed the highest Ct value (Ct ∼21), suggesting the most efficient rRNA removal, followed by Qiagen (Ct ∼19), NEB (Ct ∼16), and Thermo Fisher (Ct ∼21), which showed the lowest Ct values (Ct ∼12–13), indicating higher residual rRNA levels. Post sequencing total reads per library ranged from 90-200M reads. Post-sequencing analysis showed that the proportion of rRNA reads was less than 1% for samples processed with Vazyme and Qiagen oligo(dT) beads, 1.4% for NEB beads, and 12% for Thermo Fisher beads. Higher rRNA rates were associated with reduced data quality across several key metrics. For instance, libraries with higher rRNA levels, such as Thermo (rRNA Rate = 12.29%), showed lower exonic rate (75.78%) and elevated intronic rate (16.48%). These libraries also had higher duplication rates (38.09%) and reduced strand specificity (96.12%), indicating lower complexity in the libraries. In contrast, UHR processed with oligo dT beads from Vazyme, NEB and Qiagen with lower rRNA rates (∼0.15% - 1.5%) achieved higher exonic rates (∼85.43% – 91.76%) and lower intronic content (∼7.79% – 12.08%), reflecting more efficient transcript capture. Additionally, lower rRNA rates were associated with improved strand specificity (up to 98.5%), contributing to higher transcript detection numbers (up to 175,922) as shown in Table 1. This data also demonstrates that increased rRNA levels correlate with diminished RNA-Seq performance, reinforcing the importance of early rRNA assessment using qPCR to optimize sequencing outcomes.

## 4. Discussion

This study presents a scalable real-time 18S ribosomal RNA (rRNA) qPCR-based assay designed to assess the efficiency of rRNA depletion in libraries pre-sequencing. Since rRNA makes up most of the Total RNA and, if not effectively depleted, can dominate sequencing data, reducing the efficiency and cost-effectiveness of RNA sequencing. This study assesses the efficiency of rRNA depletion in both Total RNA and Poly A+ RNAseq libraries using a qPCR-based approach. This qPCR assay was established using 18S rRNA primers, and optimized input conditions streamlined the workflow for high-throughput sample processing to identify samples with high rRNA read percentage pre-sequencing to enable accurate transcriptome profiling.

This 18S rRNA qPCR assay can effectively screen libraries with high ribosomal RNA pre-sequencing, minimizing sequencing costs and optimizing workflow. Library dilution optimization and input normalization enabled consistent Ct values, supporting qPCR efficiency and assay reproducibility as reported by others [20, 21]. Specifically, a Ct value threshold of 16 for Total RNAseq and a Ct value of 13-14 for Poly A+ RNAseq aligns with <10% rRNA percentages in post-sequencing data (Figures 3 and 4). The use of UHR libraries as a baseline for establishing these threshold values validated the accuracy of the assay, as these controls are widely used in RNA-Seq protocol optimizations [22]. Various other studies have also proved that relative qPCR is an effective, fast, and simple method to flag problematic samples without need for complex quantifications [23].

This qPCR methodology was applied to 1,748 Total RNA and 445 Poly A+ libraries, demonstrating its effectiveness in large-scale sequencing workflows. The 18S rRNA qPCR method not only detects samples with elevated rRNA content but also can help to prioritize high-quality samples. The Ct values correlated with rRNA read percentage, with higher Ct values indicating better depletion and lower Ct values linked to higher rRNA read rates [24]. Other studies have similarly highlighted the utility of qPCR-based pre-screening in conserving sequencing depth and reducing unnecessary computational expenses associated with low-quality data [7, 12, 25].

Testing of various Oligo(dT) beads from four different vendors using the 18S qPCR assay provided valuable insights into how the assay can be used to evaluate the comparative performance of RNA-Seq kits. Previous studies show that variations in oligo(dT) beads, enzymes, and hybridization methods can significantly impact rRNA depletion and sequencing quality [9–11, 26]. Incorporating the 18S rRNA qPCR assay into experimental evaluations will allow for the identification of the most effective rRNA depletion methods and RNA-Seq kits.

The 18S rRNA qPCR Ct values closely reflect post-sequencing rRNA read percentage across all bead types, confirming effective mRNA capture especially by Vazyme and Qiagen. libraries with borderline Ct values (11–12). This method has also proved effective for streamlining and processing samples in high-throughput settings, where around 288 libraries can be processed in 8 hrs. and cost for qPCR per libraries is $2.08 which includes cost of the reagents and consumables, when run as triplicates.

## 5. Conclusion

In conclusion, this study emphasizes the significant role of the developed 18S rRNA quantitative PCR assay in effectively evaluating high rRNA presence within the RNA-Seq workflow. Its ability to detect residual rRNA allows for early identification of Library quality issues, ensuring high quality transcriptomic data. This is especially beneficial in both high-throughput and small laboratory settings, where it helps to prevent the expensive and time-consuming sequencing of poor-quality RNA. By integrating this assay, workflows become more efficient and cost-effective. This method can be used to improve RNA-Seq protocols and evaluate library kits.

## 6. Future Perspective

As RNA-seq uses continues to expand there is a need for robust quality control tools. The 18S rRNA qPCR assay described in this study offers a scalable solution to evaluate rRNA depletion efficiency early in the RNA-Seq workflow. Future applications may include integration into high throughput RNA-Seq and clinical workflows and testing of newly available kits for broader applicability. Incorporating this qPCR-based QC into RNA-Seq workflows will not only increase sequencing efficiency but also enhance reproducibility and data quality in both research and clinical settings.

## Article Highlights

● Developed a scalable qPCR-based assay targeting 18S rRNA to evaluate rRNA depletion efficiency pre-sequencing.
● Screened ∼2,200 RNA-Seq libraries (Total and Poly A+) and showed strong correlation between qPCR Ct values and rRNA read percentage post sequencing.
● Established Ct threshold value of ≥16 for Total RNA and 13-14 or greater for Poly A+ libraries.
● Evaluated Oligo-dT beads performance from four commercial vendors.
● Cost-effective and high throughput: ∼288 libraries can be processed in 8 hours at a cost of $2.08/library.

## Author contributions

Kavya Kottapalli, Hsu Chao, Harsha Doddapaneni : Conceptualization, Data Generation, Analysis and Writing.

Qi Jiang, Jimmonique Donelson, Nathanael Emerick, Sravya V Bhamidipati, Zeineen Momin, Mathew Tittu Ziad M. Khan, Korchina Viktoriya Marie-Claude Gingras : Data Generation, Analysis and Writing.

Donna Muzny and Richard Gibbs: Review & Editing.

All authors reviewed and approved the final draft of the manuscript.

## Data Availability Statement

The authors declare that the data supporting the findings of this study are available within the article and from the corresponding author upon request.

## Data Deposition

Not applicable.

## Supplemental online material

Not applicable.

## Acknowledgements

The authors are grateful to the production teams at the Human Genome Sequencing Center for data generation.

